# Immune Responses to Salmonella Typhi Antigens among Typhoid Recovered Individuals in an Endemic Region

**DOI:** 10.1101/2025.07.04.663140

**Authors:** Atharv Athavale, Adarsh Subramaniam, Aftab Hussain, Indu Rani, Deepak Kumar Rathore, Santosh Kumar Upadhyay, Anil Kumar Pandey, Praveen Kumar Malik, Amit Awasthi, Ramesh Chandra Rai

## Abstract

**Background:** Typhoid fever remains a major public health concern in endemic regions, yet the nature of antibody responses to individual Salmonella Typhi antigens and their potential role in modulating cellular immune functions are not well understood. This gap also applies to the long-term immune response after the vaccination against the disease.

**Objectives:** To evaluate the humoral and cellular immune response against the *Salmonella Typhi* antigens among the Typhoid recovered individuals.

**Methods:** ELISA was used to measure antibody responses against five *S. Typhi* antigens in plasma samples from typhoid-recovered individuals and healthy participants. Vi polysaccharide-specific plasma antibodies were tested for their ability to activate NK cells isolated from healthy donors. Flow cytometry-based assays were performed to analyze T cell and B cell profiles in the peripheral circulation of these individuals.

**Results:** Polysaccharide antigens mounted better IgA responses among typhoid-recovered individuals compared with healthy participants. Anti-*S. Typhi* IgM was significantly elevated in the typhoid group for all five antigens (p<0.05), while no significant differences were observed in IgG responses. NK cell activation assays showed comparable levels of activation markers of the NK cells isolated from healthy individuals and treated with the plasma of Typhoid recovered individuals. Analysis of B cell subsets revealed largely comparable distributions between the two groups; however, transitional B cells were elevated among the Typhoid recovered individuals. HlyE-specific memory B cells were detectable in both the groups. CD4 T cell subsets were similar between typhoid-recovered and healthy individuals, whereas differences were observed in CD8 T cell populations, with higher TEMRA cells among recovered individuals and relatively higher central memory CD8 T cells in healthy participants.

**Conclusion:** Polysaccharide antigens elicited significantly better IgA response in recovered typhoid participants, possibly reflecting mucosal immune priming from the gut. Persistent IgM antibodies after recovery suggest durable IgM responses following typhoid infection. Similar IgG levels between groups and the presence of antigen-specific memory cells in healthy individuals highlight the influence of repeated exposure in endemic settings. Elevated transitional B cells suggest the reinforcement of the B cells to the peripheral circulation during the disease which remained in the circulation of these individuals. However, elevated TEMRA population of the CD8 T cells has been observed in several other infectious diseases including this study. These findings demonstrate the complexity of humoral and cellular immune responses to *S. Typhi* in endemic regions.

## 1. Introduction

Typhoid and paratyphoid fevers, collectively known as enteric fevers, are caused by *Salmonella enterica* serovars Typhi and Paratyphi. The morbidity and mortality because of Typhoid fever is a major concern in developing countries, especially in those with poor healthcare systems. According to the World Health Organization (WHO), there are 9 million cases of typhoid annually, resulting in 1,10,000 deaths per year (1). *S.* Typhi is transmitted via the fecal-oral route, predominantly through ingestion of contaminated food and water (2). Humans are the only natural reservoirs, and chronic carriers of the bacteria and play a critical role in sustaining transmission (3). Variations in disease presentation across endemic regions and the rise of multidrug resistant strains pose challenges for timely diagnosis and effective control (2,4)

Isolating *S.* Typhi using blood culture is the gold standard method of typhoid diagnosis but it has limited sensitivity and requires adequate laboratory infrastructure (5,6). Serological tests such as Widal agglutination assay and rapid immunoassays (e.g., Typhidot, Tubex) are used in resource limited areas and typically measure antibodies against O (lipopolysaccharide/LPS) and H (flagellin) antigens. These tests suffer from low sensitivity and specificity, where the obtained results are often unreliable and show cross-reactivity with other species of Salmonella (7,8). There is no reliable method for the detection of chronic typhoid and serological tests may not distinguish between current and past infection (1).

There is a gap in our understanding about *S.* Typhi infection and its pathogenesis, where the human-restricted nature of this pathogen has limited our ability to model host-pathogen interactions effectively. Adaptive immunity against *S.* Typhi includes both cellular and humoral arms (9). Humoral responses involve antibodies to diverse antigens such as surface polysaccharides (LPS [O antigen], Vi capsule) and bacterial proteins (e.g., flagellin [H antigen], HlyE toxin, outer membrane proteins). The role of antibodies in protection against *S.* Typhi is not fully understood, however anti-Vi-polysaccharide responses correlate with vaccine-induced immunity (9,10). Recent studies have identified IgA against HlyE and LPS as strong markers of acute typhoid infection (11). However, little is known about how long these antibodies persist post-infection or how they differ among individuals of endemic area without clinical illness. In addition to antibody responses, the phenotypic landscape of circulating B cell and T cell subsets among typhoid-recovered individuals remains poorly characterized, particularly in endemic regions. Understanding how infection shapes memory B cell and T cell compartments is important for interpreting long-term immune responses and may provide insights into vaccine-induced immunity against S. Typhi.

Key virulence determinants of *S.* Typhi include the Vi capsular polysaccharide, Type III secretion systems (T3SS-1 and T3SS-2), and typhoid toxin, which facilitate intestinal invasion, intracellular survival within macrophages, and systemic dissemination (12). Protective immunity against typhoid fever involves antibodies directed against the Vi antigen, robust Th1 responses characterized by IL-12 and IFN-γ production, and cytotoxic T cell lymphocyte activity that promotes intracellular killing of bacteria (13,14). A detailed understanding of these correlates of protection underpins current vaccine strategies (15). Both B cells and T cells play important roles in shaping protective immunity against intracellular bacterial pathogens. B cells contribute to the generation of antigen-specific antibody responses and memory B cell populations, while T cells mediate cellular immunity through helper and cytotoxic responses that facilitate bacterial clearance. However, information regarding the distribution of circulating B cell and T cell subsets following recovery from typhoid infection remains limited.

*S.* Typhi capsule consists of Vi polysaccharide and is encoded by the genes of Salmonella Pathogenicity Islands (SPI) (3,16). It has been noted in a study from India, that about 10% of the fresh isolates lack Vi-polysaccharide antigens (17). However, strains of *S.* Typhi with Vi polysaccharide are more infectious and virulent than Vi-negative strains (2). H-protein, a flagellar antigen recognized by Toll-like receptor 5 (TLR5), triggers strong antibody responses, while the Vi capsular polysaccharide antigen acts as a virulence factor by masking O antigens and protecting the bacteria from complement-mediated lysis and phagocytosis (18,19). LPS, composed of lipid A, core oligosaccharide, and O-specific polysaccharide chains, is recognized by TLR4 and induces robust inflammatory responses (20,21). Interactions among these antigens are complex; for instance, Vi expression can suppress immune recognition of LPS, altering host-pathogen interactions (22–24). In addition to these antigens, Outer Membrane Proteins (OMPs) represent significant immunogenic component involved with host-pathogen interactions during *S.* Typhi infection. These proteins play essential roles in the pathogen’s adaptation to environmental conditions, motility, adherence, and host cell colonization. *S.* typhi OMPs have demonstrated potent immunogenic properties, eliciting long-lasting and protective immunity, which makes them a potential vaccine candidate, and diagnostic antigen.

In endemic regions, where typhoid fever is widespread, immune responses to *S.* Typhi are complicated by frequent exposure to the pathogen. How different antigenic components contribute to long-term immunity in endemic settings, where individuals may experience repeated exposure and subclinical infections, remains poorly understood.

This study focuses on evaluating the antibody responses-IgA, IgG, and IgM; against five critical *S.* Typhi antigens: Vi polysaccharide, H-protein, LPS, Hemolysin E and Outer Membrane Protein and examining the distribution of circulating B cell and T cell subsets to understand the long-term immune landscape shaped by *S.* Typhi exposure in endemic regions. Plasma samples were collected three months after typhoid to assess the dynamics of the humoral immunity, with a focus on identifying immunological patterns that could inform vaccine response studies. To further improve our understanding about cellular innate immunity, we tested whether Vi-polysaccharide specific antibodies can activate the NK cells, owing to the role of the NK cells in eliminating cells infected by an intracellular pathogen. In addition, we characterized circulating B cell and T cell subsets in order to understand the broader immune landscape following recovery from typhoid infection. The findings of this study reveal heterogeneous humoral responses among typhoid-recovered individuals against these antigens. Notably, the substantial immune responses observed in healthy participants also highlight the *S.* Typhi exposure in this region, emphasizing the challenges posed by the pathogen endemicity.

## 2. Materials and Methods

### 2.1 Ethics statement

This study was conducted at the BRIC-Translational Health Science and Technology Institute, Faridabad, India. Participant enrolment and sample collection were done at ESIC Medical College and Hospital, Faridabad. Written informed assent/consent was obtained from each of the study participants. This study was approved by the Institutional Ethics Committees from both institutes [*ESIC Hospital and Medical College, Faridabad File no. 134 X/11/13/2022-IEC/110 and THSTI Faridabad Ref No: THS 1.8.1/* (*152*)].

### 2.2 Participant details

Blood samples were collected from a total of 80 participants **(Table 1)**. Their age, gender, infection date (in the case of typhoid-recovered patients) and blood collection date were recorded. The study population consisted primarily of rural residents and healthcare workers.. Among the 80 participants we recruited for the study, 33 had recovered from typhoid fever at least for 3 months, while 47 were healthy participants and had no recorded history of the disease.

**Table 1:** Participant demographics.

|  |  | **Typhoid Recovered** | **Healthy** |
| --- | --- | --- | --- |
| Sex | Male | 15 | 27 |
|  | Female | 18 | 20 |
| Age (years) | Range | 13-53 | 12-58 |
|  | Median | 25 | 24 |
| Months since infection | Range | 3-10 | - |
|  | Median | 3 | - |

### 2.3 Inclusion and Exclusion criteria

The study included participants ages 18–59 years tested for Typhoid fever. Eligible individuals were either healthy or were recovered from typhoid fever at least three months prior to blood collection. A positive Widal test was not considered as a confirmatory test; a positive Typhidot IgM/IgG or blood culture tests were used instead. Pregnant women and people with immunocompromising comorbidities were excluded from the study.

### 2.4 Blood collection and sample processing

Eight ml of blood was collected from each participant in the Greiner Bio-One tubes (Cat. No. 22-040-134) coated with sodium heparin. Blood samples were processed immediately to isolate plasma and peripheral blood mononuclear cells (PBMCs) using Lymphoprep density gradient centrifugation method. Plasma was stored at -80 °C and PBMCs were resuspended in fetal bovine serum (FBS) supplemented with 10% dimethyl sulfoxide (DMSO) and then stored in liquid nitrogen (25).

### 2.5 ELISA

#### 2.5.1 Polysaccharide antigens

Anti- *S.* Typhi IgA, IgG and IgM antibody responses are measured against Vi polysaccharide and Lipopolysaccharide after the three-months from infection of the typhoid patients by using indirect ELISA. Nunc Maxisorp 96-well plate was coated with 100 µL of poly-l-lysine (2 µg/ml) and incubated at 37°C for 2 hours. The poly-l-lysine coating was not done in case of the H antigen. All antigens were diluted in sterile PBS (2 µg/ml for Vi polysaccharide; 1 µg/ml for Lipopolysaccharide and H-protein). Antigens were coated and incubated overnight (37°C for Vi polysaccharide and lipopolysaccharide; 4°C for H-protein). Coated plates were blocked with a 200 µl block buffer (1% BSA in PBS) and incubated for 1 hour at 37°C. Plasma samples were diluted 1:100 in a diluent (1% BSA, 0.05% Tween-20 in PBS) and 100 µl/well was added and incubated for one hour at 37°C. Then, 1/10000 secondary antibody was diluted in a buffer containing 1% BSA, and 0.05% Tween-20 in PBS and 100 µl was added to each well and incubated. After an hour 100 µl pNPP (p-Nitrophenyl Phosphate) substrate is added to each well which is diluted in Tris-MgCl_2_ and incubated for an hour at room temperature in dark conditions. At last, 50 µl 3M NaOH is added to stop the reaction. After every step of ELISA, 4 to 5 washing is given with 0.05% Tween-20 in PBS except after the addition of substrate. Then, optical density is measured using a plate reader at 405 nm and 490 nm. The readings at 490 nm were subtracted from those at 405 nm to correct for optical imperfections.

#### 2.5.2 Protein Antigens

For the protein antigens (H-protein, Hemolysin E and OMP), similar ELISA was performed with some modifications. HRP-TMB was used as the conjugate-substrate combination instead of alkaline phosphatase-pNPP; because it was found to give better and consistent results during the optimization experiments. Hence, the stop solution was changed to 2N H_2_SO_4._

### 2.6 Avidity assay

Antibody avidity against S. Typhi Vi polysaccharide and LPS was measured using a modified indirect ELISA protocol as described above. Briefly, plates were coated with poly-L-lysine (10 µg/mL) and incubated at 37 °C for 2 hours, followed by coating with Vi antigen or LPS (0.5 µg/mL) overnight at 37 °C. Plates were blocked with 1% BSA in PBS and plasma samples diluted in sample diluent (1% BSA + 0.1% Brij-35 in PBS) were added and incubated for 1 hour at 37 °C.

After sample incubation, 8 M urea (100 µL/well) was added to half of the replicate wells, while wash buffer was added to the remaining wells as control, and the plates were incubated for 10 minutes at 37 °C. Plates were then washed and incubated with alkaline phosphatase–conjugated secondary antibody (1:10,000 dilution) for 30 minutes at 37 °C. The reaction was developed using pNPP substrate and stopped with 3 M NaOH. Optical density was measured at 405 nm and 490 nm using a microplate reader, and background correction was performed by subtracting the OD at 490 nm from the OD at 405 nm.

### 2.7 Antibody-dependent NK cell activation

Nunc MaxiSorp® flat-bottom plate was coated with streptavidin (5 µg/ml) and incubated overnight at 37°C. After 3 washes with PBS (200 µl/well), biotinylated Vi-polysaccharide (26,27) (3 µg/ml) was added, and the plate was incubated at 37°C for 3 hours. After 5 washes with PBS, diluted plasma (plasma: PBS; 1:10) from typhoid-recovered patients was added and the plate was incubated at 37°C for 2 hours. NK cells obtained using Miltenyi MACS NK cell isolation kit (cat no. 130-092-657) were added to the plates after 5 PBS washes. 200 µl complete RPMI media was added, and the plate was incubated at 37°C for 12 hours. The cells from the microplate were carefully scraped off with 100 µl/well FACS buffer (PBS + 1% FBS) and transferred to a 96-well round bottom plate using a multichannel pipette. The step was repeated if required, to get most of the residual cells. The cells were washed twice by adding a FACS buffer and centrifuged at 3000 rpm for 5 minutes. Excess FACS buffer was decanted carefully, and a cocktail of antibodies was added (40 µl/well) **(Table 2)**. Samples were acquired at BDFACSCanto II and analyzed using FlowJo software (version v10). The gating strategy of the data analysis has been shown in **Supplementary Figure 1**.

**Table 2:**
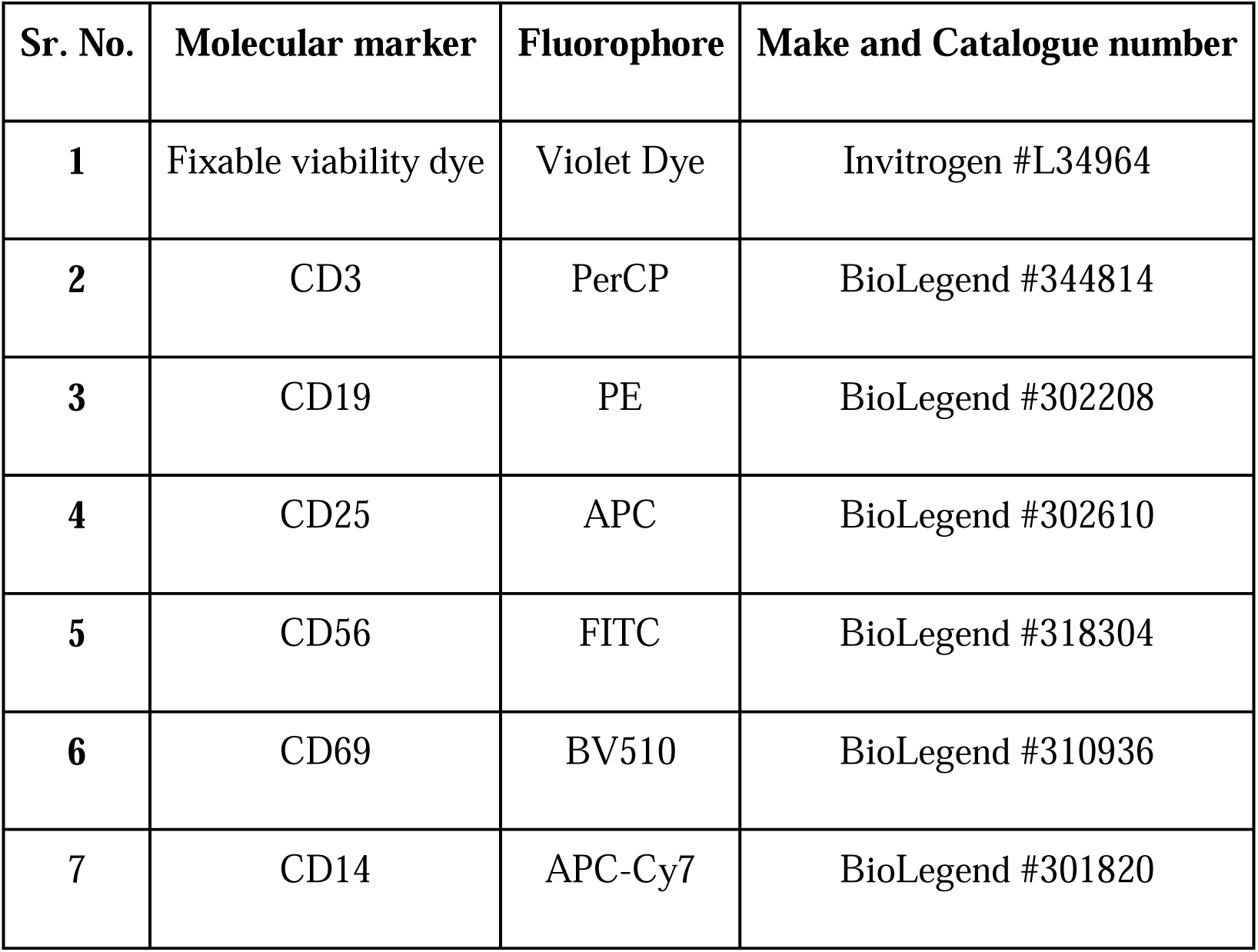
Molecular markers used for estimation of NK cell proliferation and activation.

| Sr. No. | Molecular marker | Fluorophore | Make and Catalogue number |
| --- | --- | --- | --- |
| 1 | Fixable viability dye | Violet Dye | Invitrogen #L34964 |
| 2 | CD3 | PerCP | BioLegend #344814 |
| 3 | CD19 | PE | BioLegend #302208 |
| 4 | CD25 | APC | BioLegend #302610 |
| 5 | CD56 | FITC | BioLegend #318304 |
| 6 | CD69 | BV510 | BioLegend #310936 |
| 7 | CD14 | APC-Cy7 | BioLegend #301820 |

### 2.8 Labelling of HlyE Protein with the Alexa Fluor-488

Labelling of the HlyE protein was performed using the Alexa Fluor Microscale Protein Labelling kit (Cat No. #A30006; Thermo Fisher Scientific) as per the manufacturer’s instructions. Fluorophore-labelled HlyE was aliquoted and stored at −20 °C for further use in testing antigen-specific memory B cell response.

### 2.9 T cell and B cell phenotyping using flow cytometry

Peripheral blood mononuclear cells (PBMCs) isolated from study participants were used for immunophenotyping of B cell and T cell subsets (Table 3). Cryopreserved PBMCs were thawed, washed with RPMI medium supplemented with fetal bovine serum and resuspended in FACS buffer (PBS + 1% FBS). Cells were stained with fluorochrome-conjugated antibodies against CD3, CD4, CD8, CD19, CD27, CD38, IgD, CD24, CD45RA and CCR7. Surface staining was performed for 30 minutes at 4°C in the dark. After washing with FACS buffer, cells were acquired on a BD FACSymphony A3 flow cytometer and analyzed using FlowJo software (version 10.10.0).

**Table 3:**
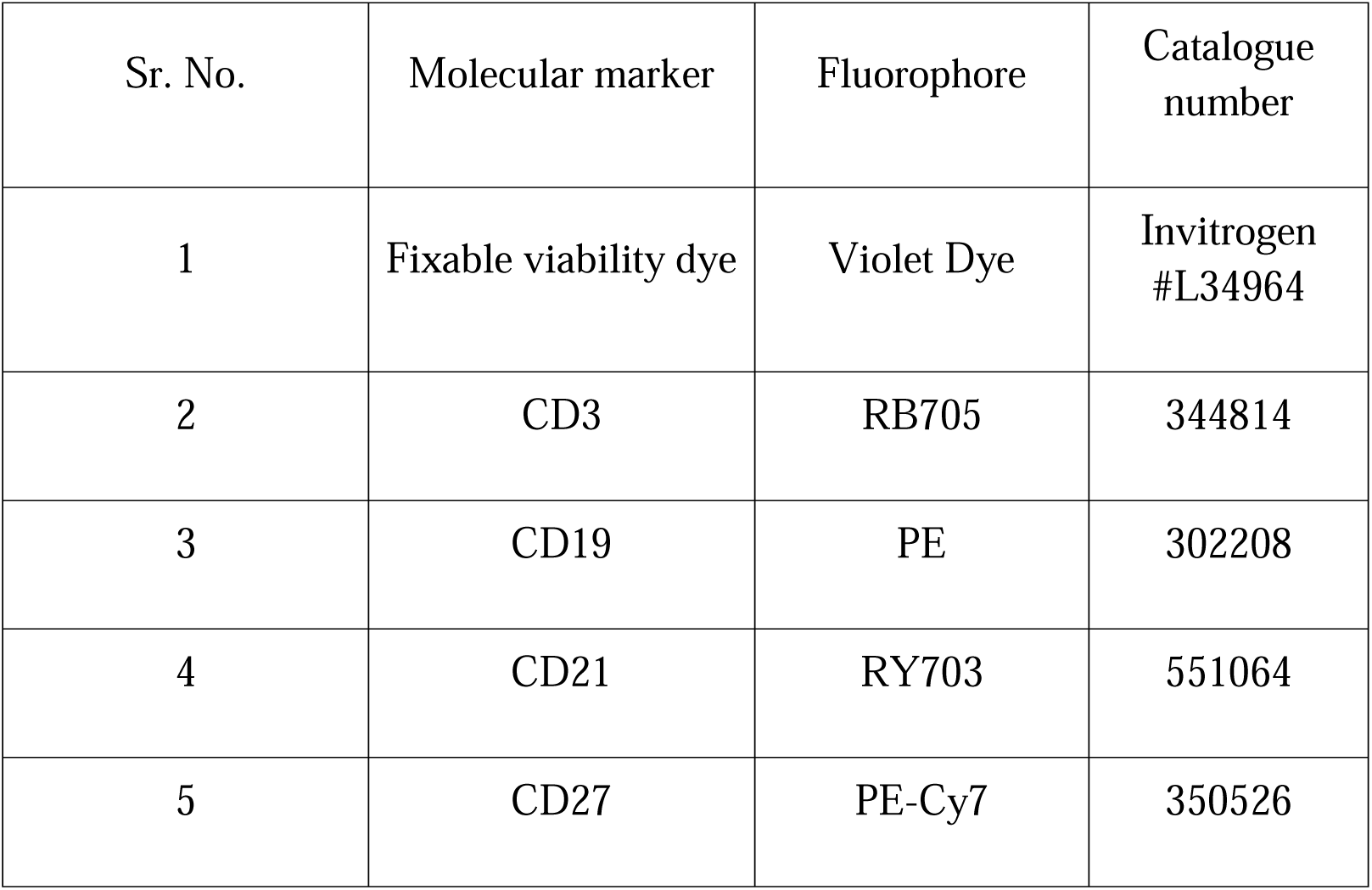
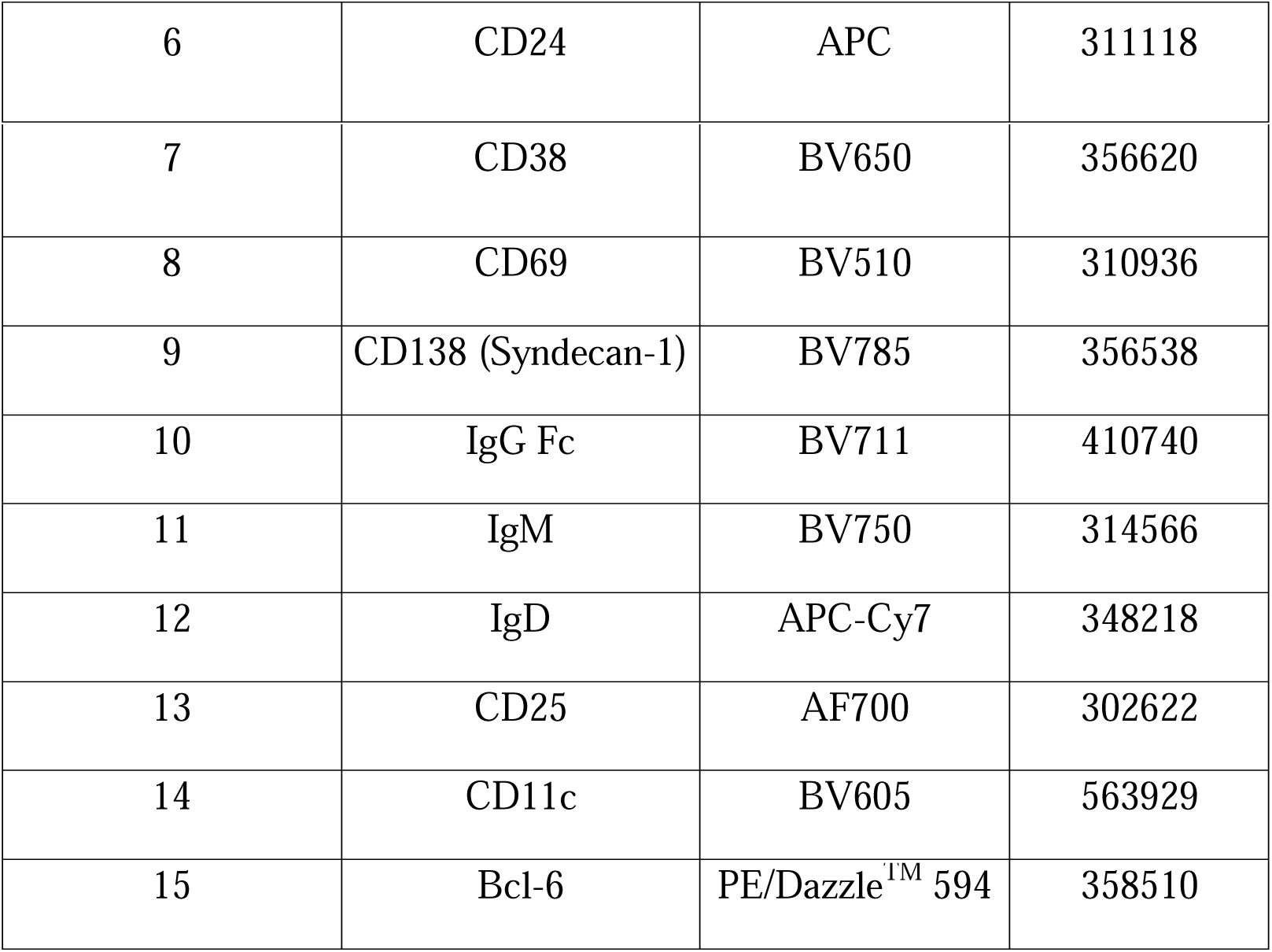
Molecular markers used for analyzing B cell and T cell phenotyping.

B cells were identified as CD3 CD19 cells. Naïve (CD27 IgD), switched memory (CD27 IgD), unswitched memory (CD27 IgD) and double negative (CD27 IgD) B cell populations were analyzed within the B cell compartment. Classic memory B cells were defined as CD27 CD38 , and transitional B cells were defined as CD27 CD24 CD38-low.

T cells were gated as CD3 lymphocytes, followed by separation into CD4 and CD8 populations. T cell subsets were defined based on CD45RA and CCR7 expression as naïve (CD45RA CCR7), central memory (CD45RA CCR7), effector memory (CD45RA CCR7) and TEMRA (CD45RA CCR7) cells.

### 2.10 Statistics

All the data visualization and statistical analyses were done using GraphPad Prism 10.4.2. Outliers were removed using the ROUT method (Q=1%). All the datasets were checked for normal distribution first. The Mann-Whitney test was used for statistical analysis to compare the two datasets. p value <0.05 was considered statistically significant.

## 3. Results

### 3.1 Polysaccharide antigens lead to a better IgA response

Anti-polysaccharide IgA (to LPS and Vi) was significantly higher in recovered participants than in healthy ones. For anti-LPS IgA, median OD was ∼0.5852 in typhoid-recovered vs 0.4064 in healthy participants (p=0.0087). For anti-Vi IgA, median OD was ∼0.406 vs 0.273 (p=0.0140) **[Fig. 1]**. IgA responses to H antigen, and OMP did not differ significantly; however, IgA response against HlyE showed significant difference among Typhoid recovered and healthy individuals **[Fig. 1]**.

**Fig. 1.**
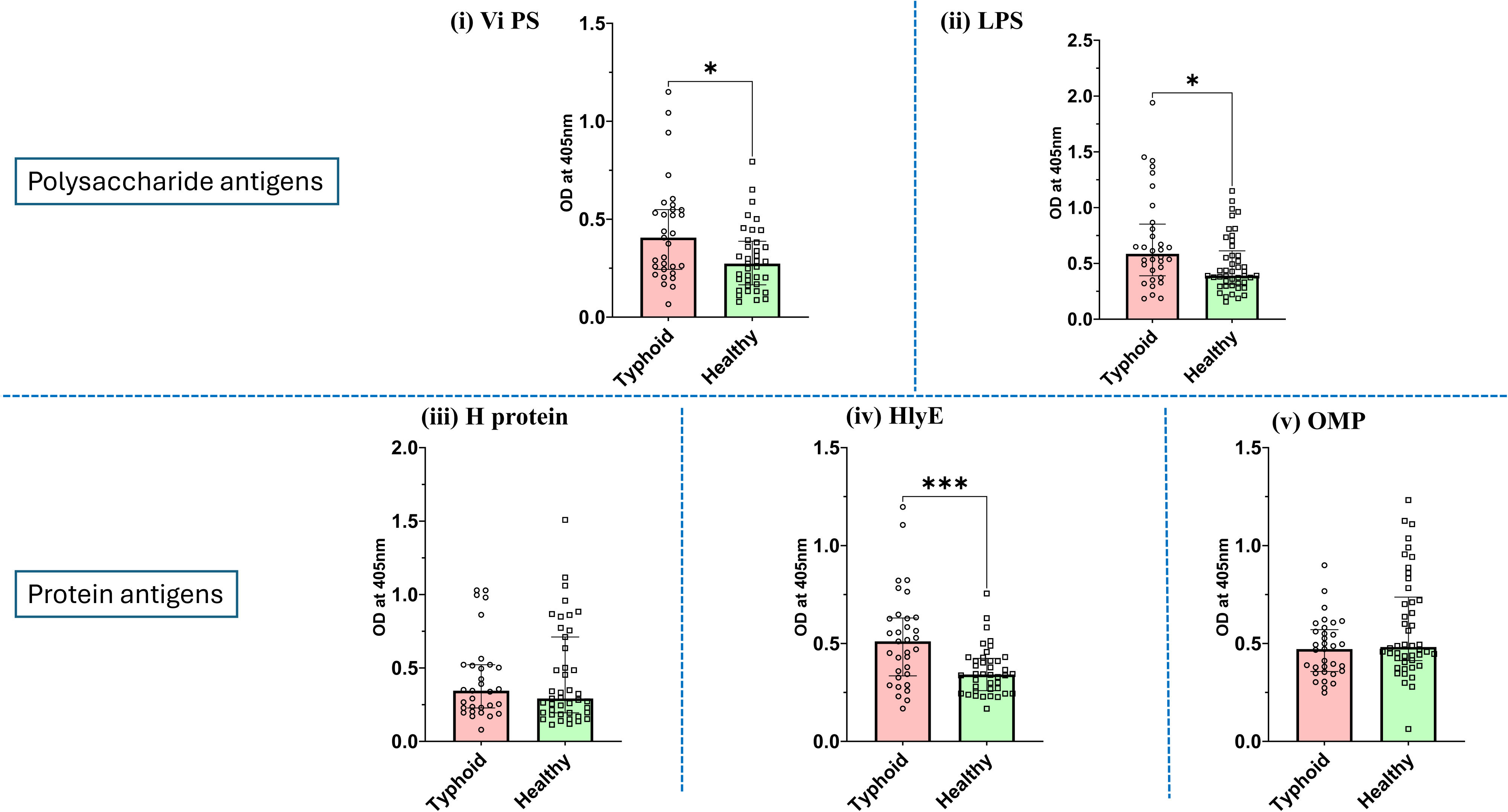
IgA ELISA against Vi PS, LPS, H protein, HlyE and OMP

### 3.2 Sustained IgM response against *S.* Typhi antigen among study participants

We observed sustained IgM response against the S. Typhi antigens among the study participants. However, IgM levels were significantly elevated in the typhoid-recovered group for all five antigens **[Fig. 2]**. IgM to H antigen, HlyE, and OMP were all roughly twofold higher in the typhoid group (all p<0.01). This elevated IgM level among typhoid recovered individuals suggests durability of IgM response after typhoid, complicating the diagnosis in endemic regions. Notably, healthy participants samples showed detectable IgA and IgM levels against all the antigens, indicating that they may have been exposed to the pathogen and remained asymptomatic.

**Fig. 2.**
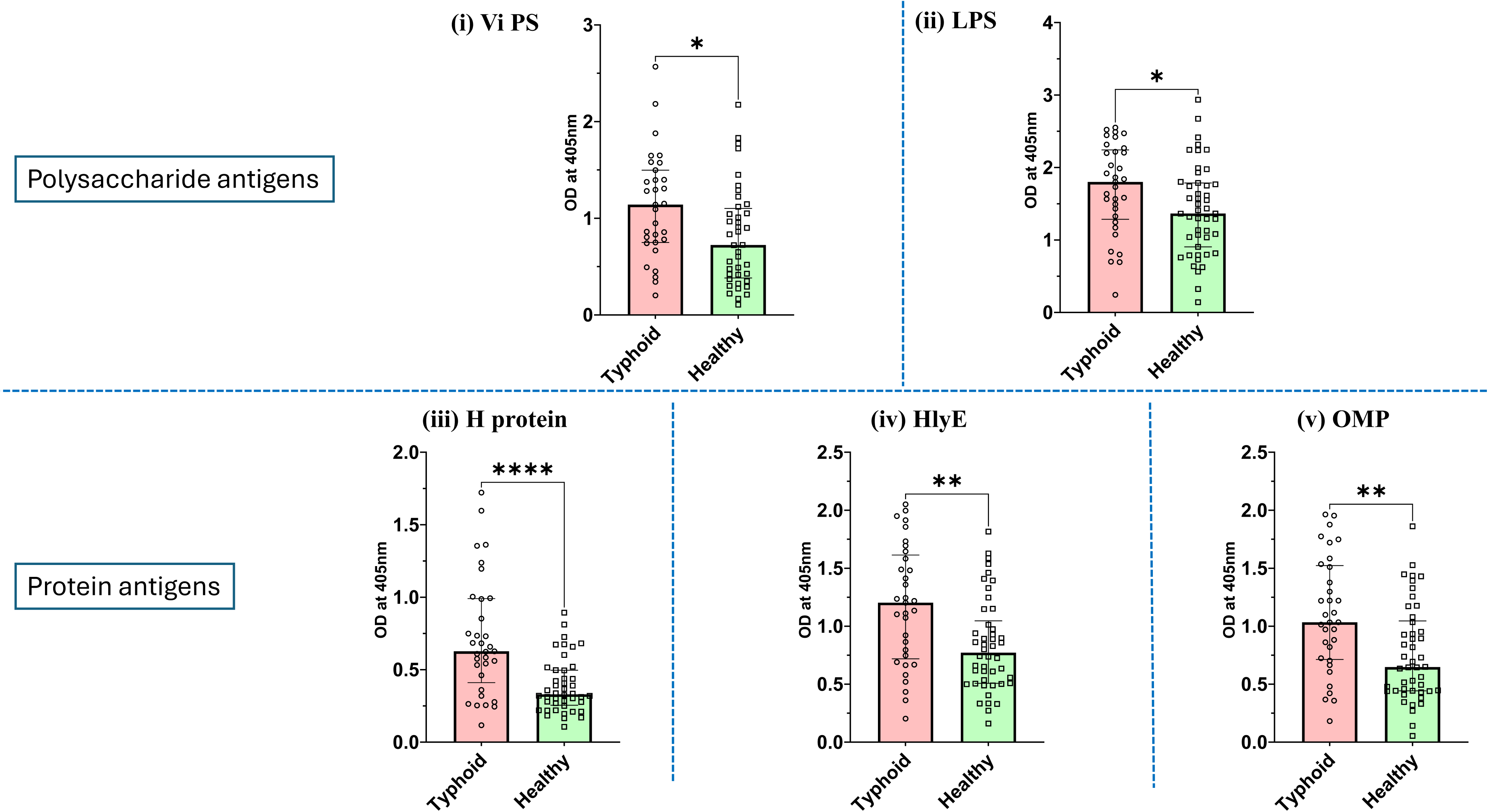
IgM ELISA against Vi PS, LPS, H protein, HlyE and OMP

### 3.3 IgG antibody responses and IgG antibody avidity (against Vi PS) did not differ between typhoid-recovered and healthy individuals

No significant differences were observed in antigen-specific IgG levels between the groups. Mean IgG ODs were similar in recovered vs healthy participants for all antigens (p>0.1). This indicates that by >3 months post-infection, IgG levels against these antigens had declined to the background levels of the endemic population **[Fig. 3]**. The chaotropic agent-based modified ELISA assay for testing the avidity of the IgG antibodies yielded interesting results. The median IgG antibody avidity index for both participant groups exceeded 70%, substantiating the presence of high-affinity antibodies targeting the Vi polysaccharide antigen **[Fig. 3(i)(b)]**. A summary of the serological data has been provided in **Table 4**.

**Fig. 3.**
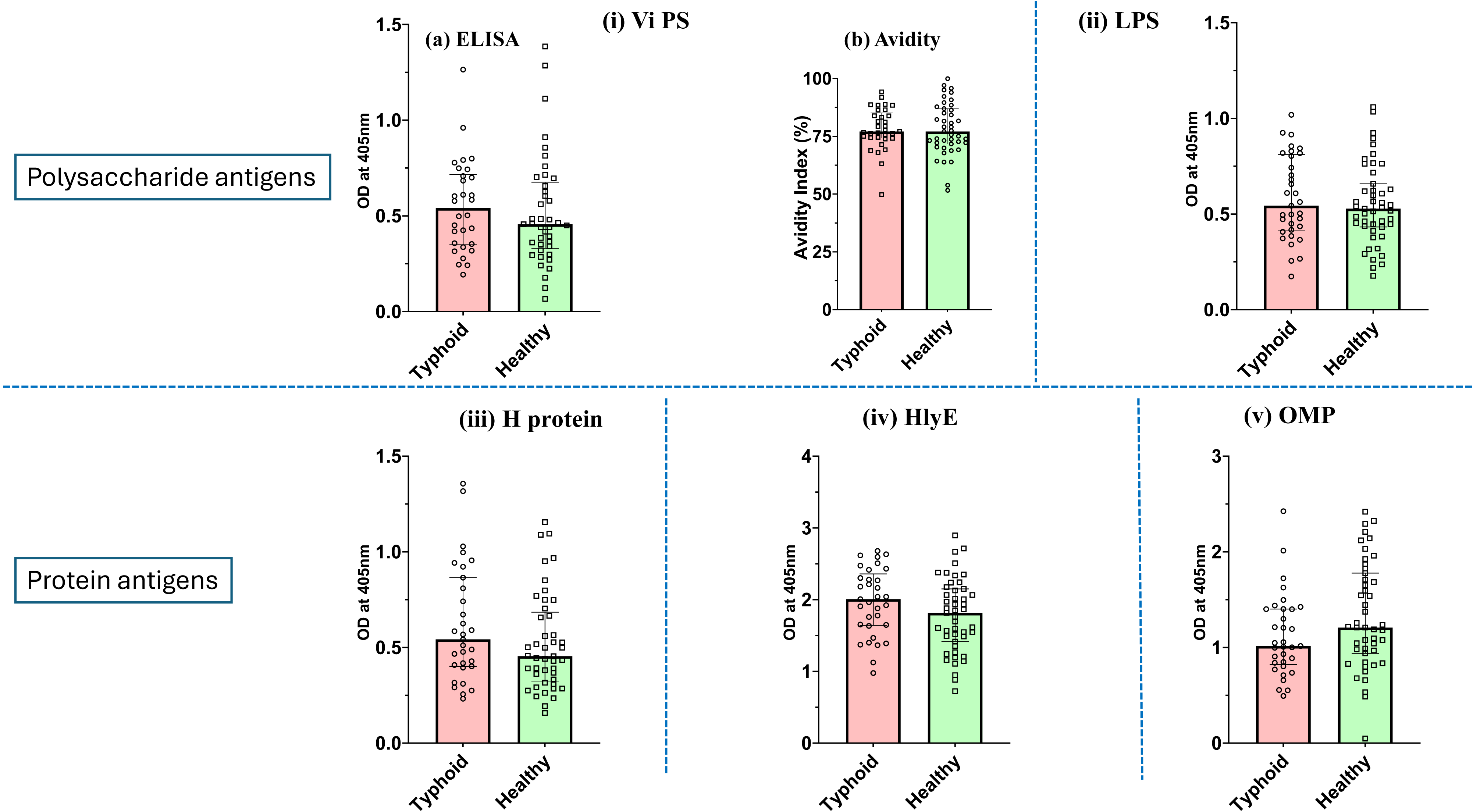
IgG ELISA against Vi PS, LPS, H protein, HlyE and OMP; IgG Avidity against Vi PS

**Table 4:**
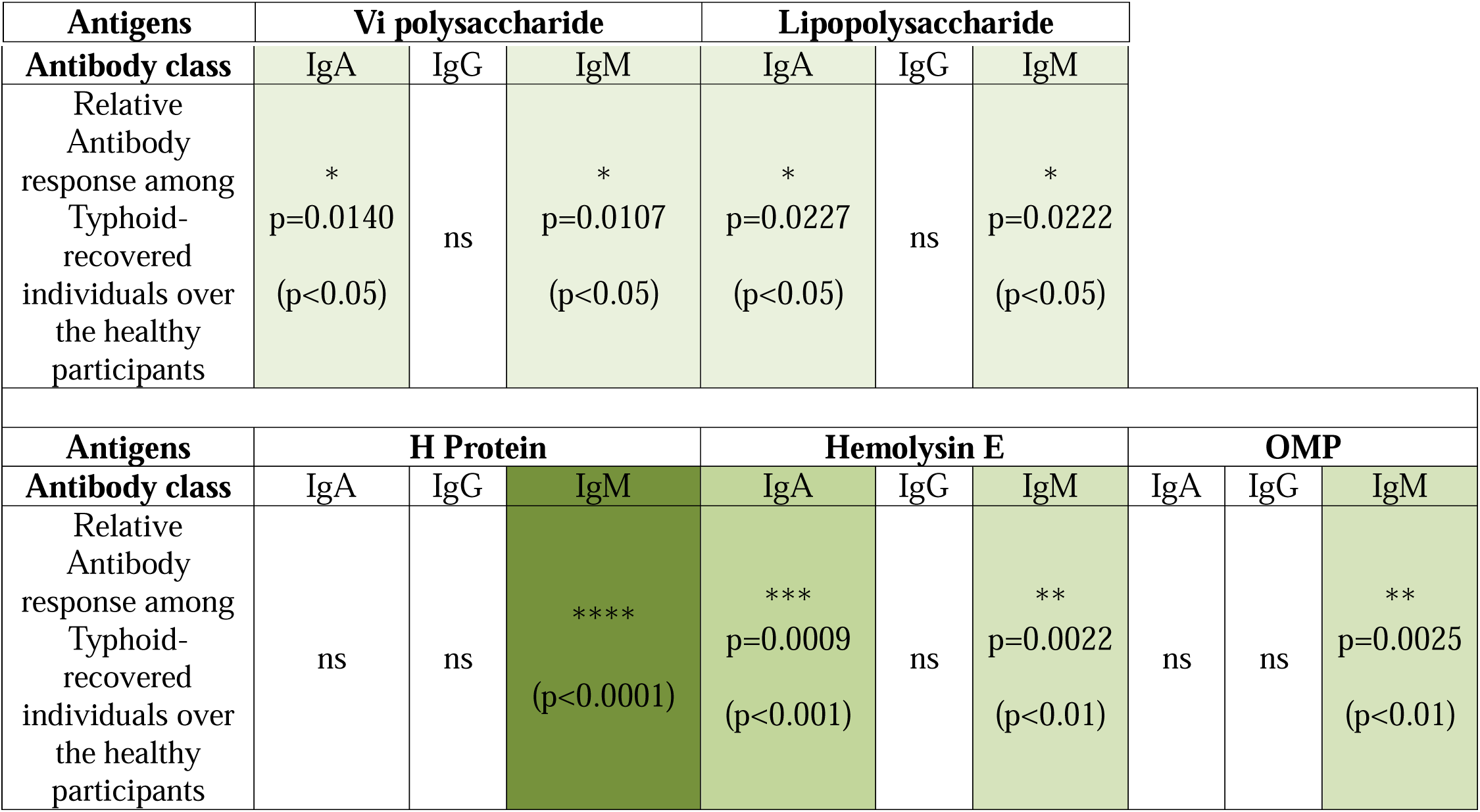
Antigen-wise seropositivity among individuals after 3 months of recovery from typhoid (ELISA)

### 3.4 No sex-specific disparities observed in serological immune responses

The study cohort demonstrated a balanced representation of both sexes, as evident from **Table 1**. Specifically, the participant groups included approximately 55% females among typhoid-recovered individuals and 43% females in the healthy group. Sex-disaggregated analysis of the immune responses was performed to evaluate any potential disparities between immune responses in males and females against *S.* Typhi. Our analysis revealed that there is no statistically significant difference in serological immune responses between these participants **[Supplementary Figures 2-6]**. This data indicates that sex does not appear to influence the immune response at recovery to typhoid infection.

### 3.5 NK cell response to antibody-dependent activation

We investigated the potential of anti-Vi polysaccharide (Vi-PS) antibodies present in the plasma samples of typhoid-recovered individuals to be able to activate natural killer (NK) cells isolated from the PBMCs of the healthy participants. The primary hypothesis was that the NK cells isolated from healthy participants would exhibit distinct immune responses when exposed to anti-Vi PS antibodies. To test this, we measured NK cell activation and proliferation using molecular markers CD16, CD25 and CD69 after incubating NK cells with typhoid-recovered participant’s plasma and culturing the cells for 12 hours. Cells incubated with plasma showed similar levels of activation and proliferation markers as those cultured without any plasma **[Fig. 4]**. Probably, due to the endemicity of the disease in India, the control samples also behaved like the plasma treated ones. Another possibility is that the interaction of the antigen with antibodies present in the plasma of the typhoid participants is needed to activate the NK cells.

**Fig. 4.**
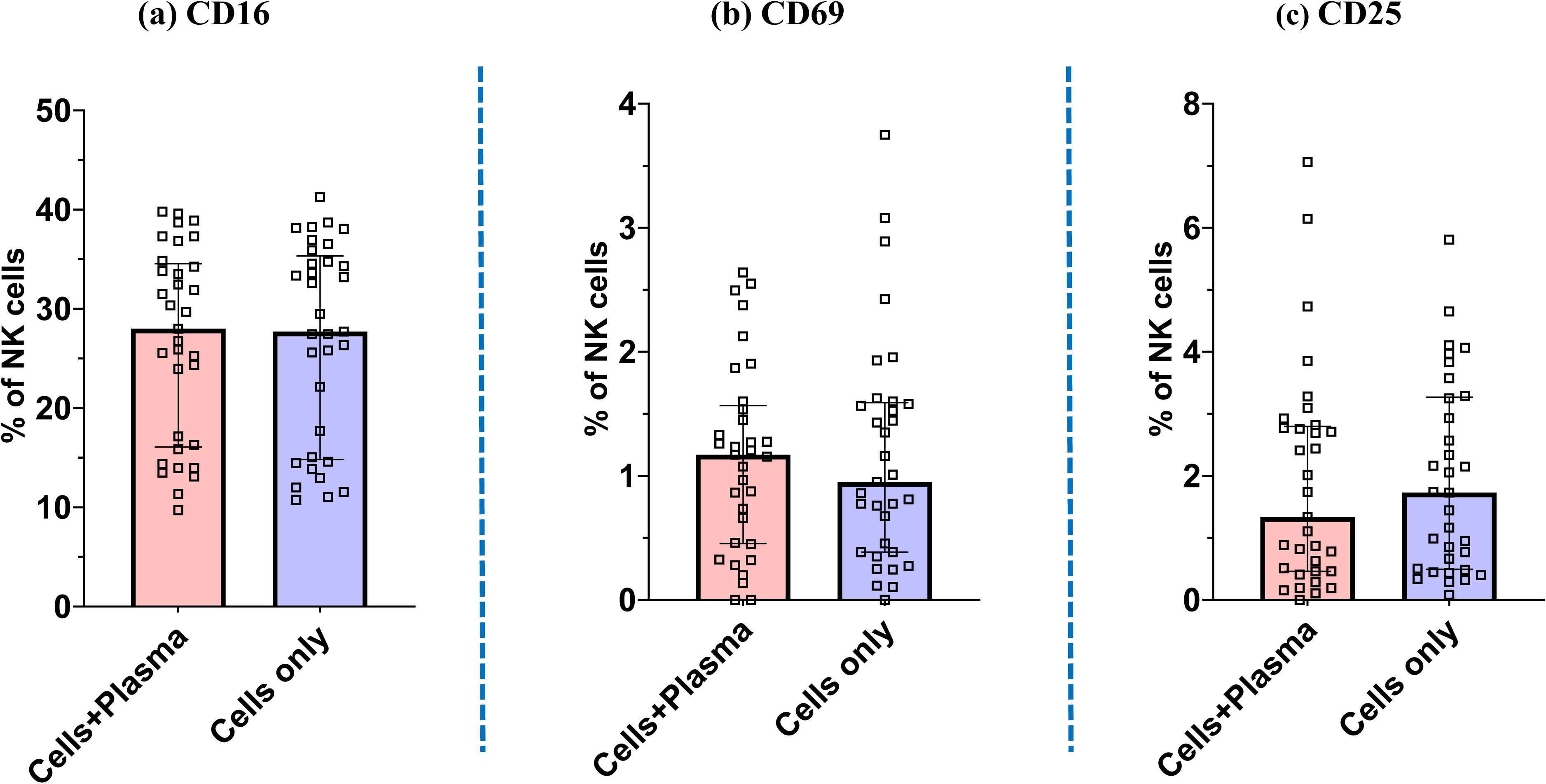
Percentage of NK cell proliferation and activation markers after antibody-dependent activation

**Fig. 5.**
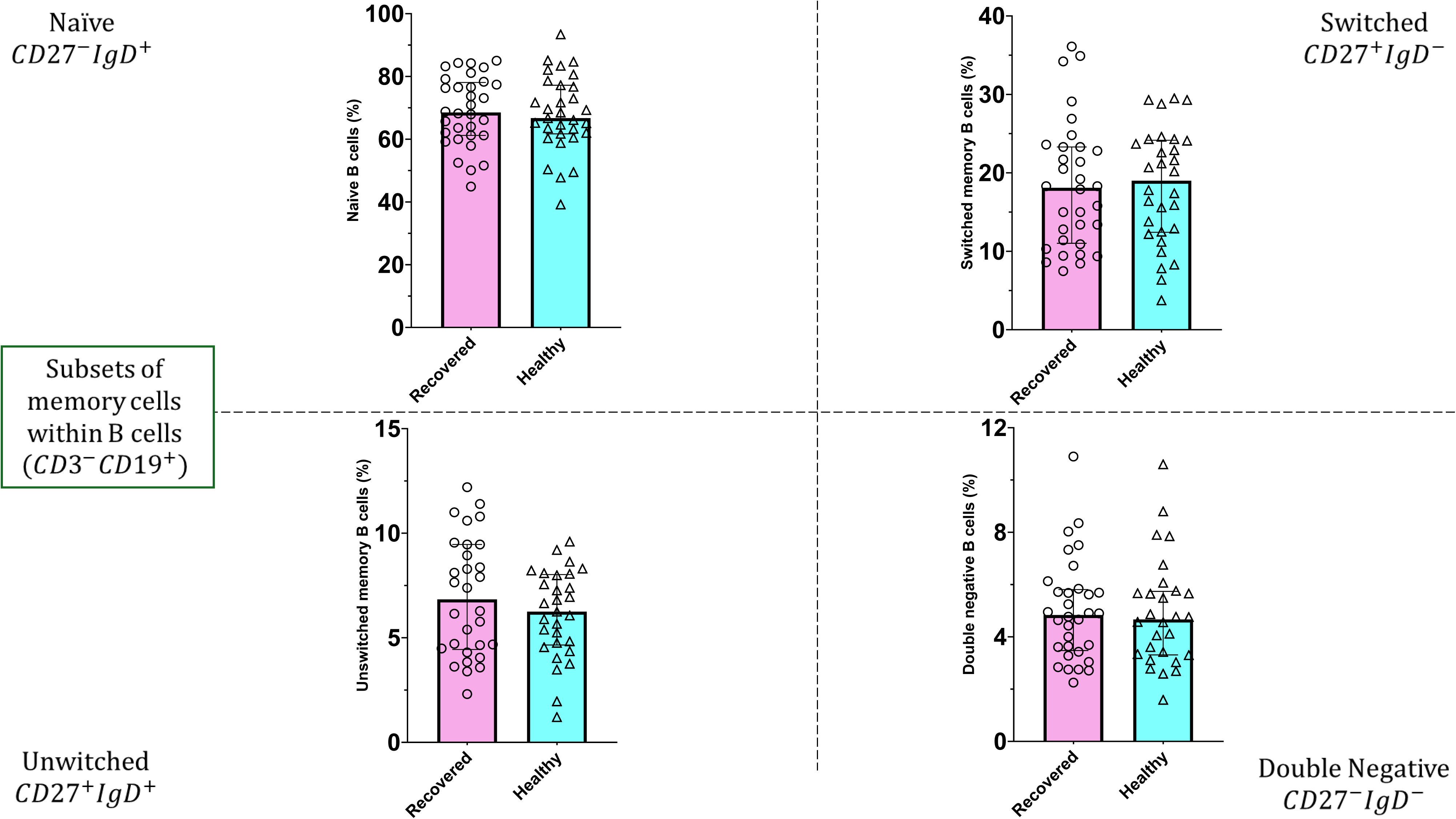
Distribution of B cell subsets within CD3 CD19 B cells showing naïve (CD27 IgD), switched memory (CD27 IgD), unswitched memory (CD27 IgD) and double negative (CD27 IgD) populations.

### 3.6 Distribution of circulating B cell and T cell subsets

To further understand the immune landscape following typhoid infection, circulating B cell and T cell subsets were analyzed using flow cytometry.

Within the B cell compartment, the proportions of naïve, switched memory, unswitched memory and double negative B cells were comparable between typhoid-recovered and healthy individuals **(Fig. 6)**. Similarly, the frequency of classic memory B cells (CD27 CD38) did not differ substantially between the two groups. However, a modest but statistically significant difference was observed in transitional B cells (CD27 CD24 CD38-low), where typhoid-recovered individuals showed slightly higher frequencies compared with healthy participants.

**Fig. 6.**
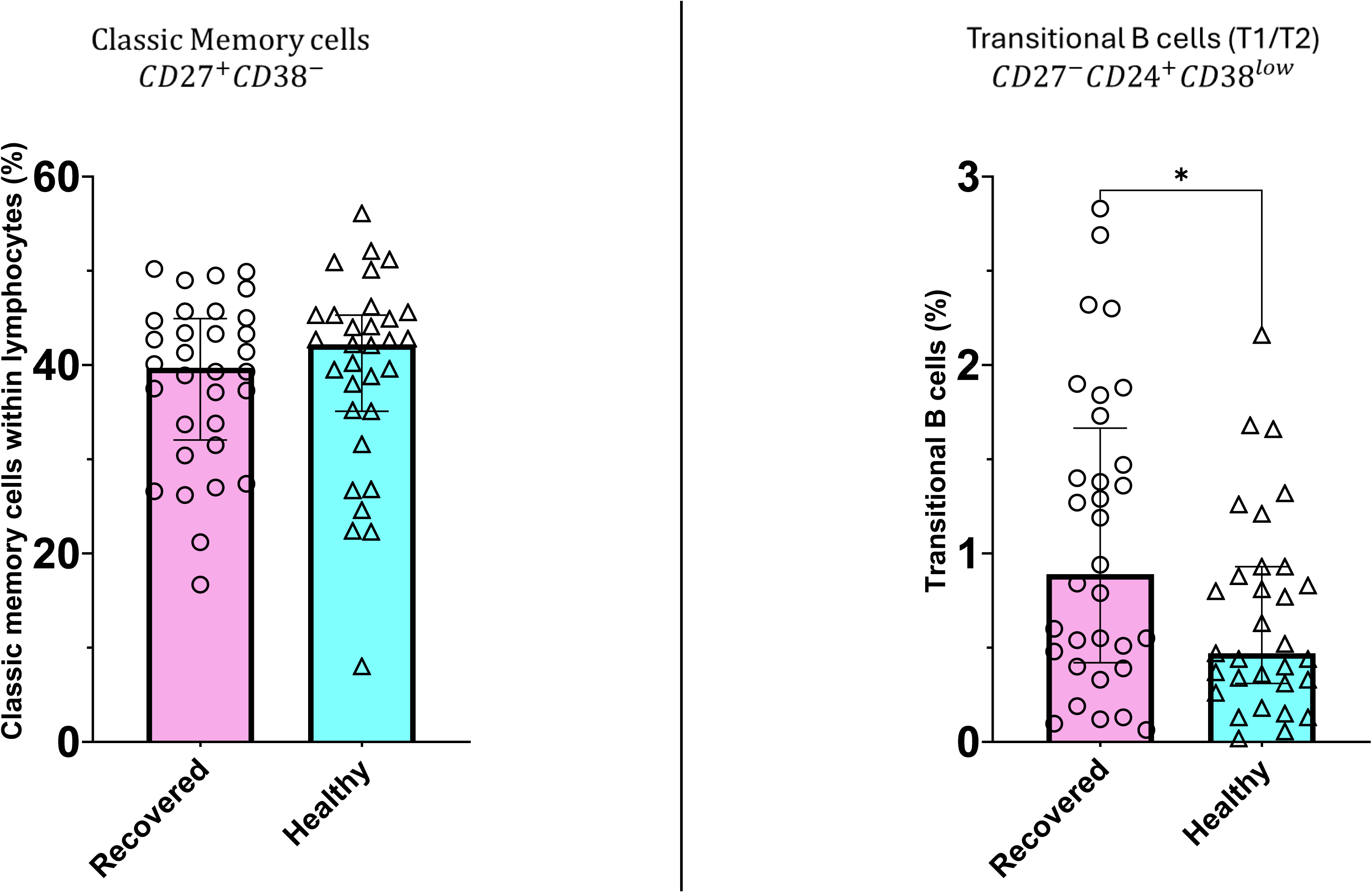
Frequency of classic memory B cells (CD27 CD38) and transitional B cells (CD27 CD24 CD38-low) among circulating lymphocytes.

We also evaluated antigen-specific memory B cells in the two study groups; using HlyE as it elicited better humoral response among protein antigens. HlyE-specific memory B cells were detectable in both typhoid-recovered and healthy participants, indicating prior exposure to the antigen in this endemic setting. The overall frequency of HlyE-specific memory B cells and the proportion of IgG-producing or IgM-producing cells within this population did not differ significantly between the two groups **(Fig. 7)**.

**Fig. 7.**
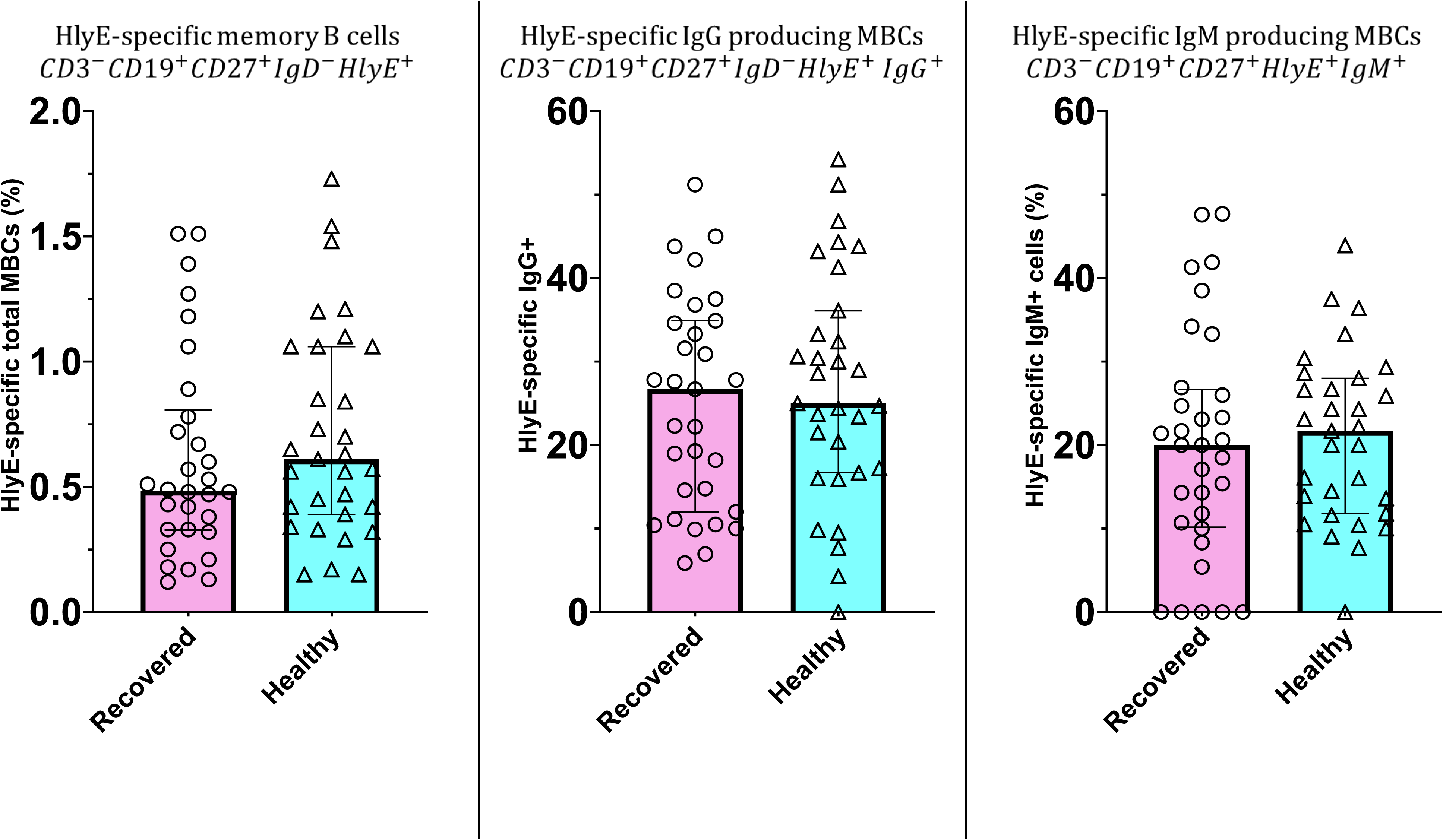
Frequency of HlyE-specific memory B cells and the proportion of IgG and IgM cells within this population.

Analysis of T cell subsets revealed that the distribution of CD4 T cell populations (naïve, central memory, effector memory and TEMRA cells) was largely similar between typhoid-recovered and healthy individuals **(Fig. 8)**. On the other hand, differences were observed within the CD8 T cell compartment. Typhoid-recovered individuals showed higher frequencies of CD8 TEMRA cells, while healthy participants showed relatively higher levels of CD8 central memory T cells **(Fig. 9)**. These observations suggest alterations in CD8 T cell differentiation and or distribution following typhoid infection.

**Fig. 8.**
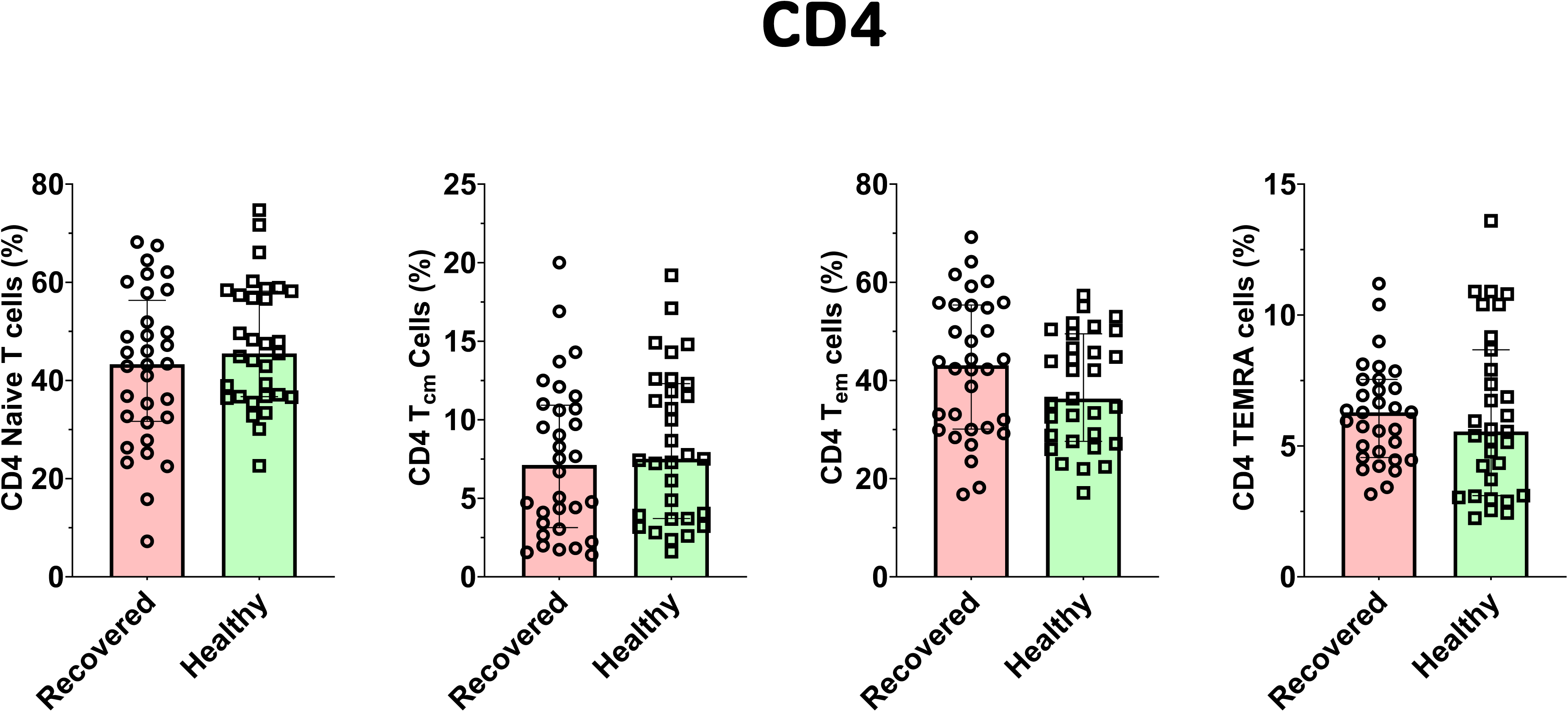
Distribution of CD4 T cell subsets (naïve, central memory, effector memory and TEMRA) among typhoid-recovered and healthy participants.

**Fig. 9.**
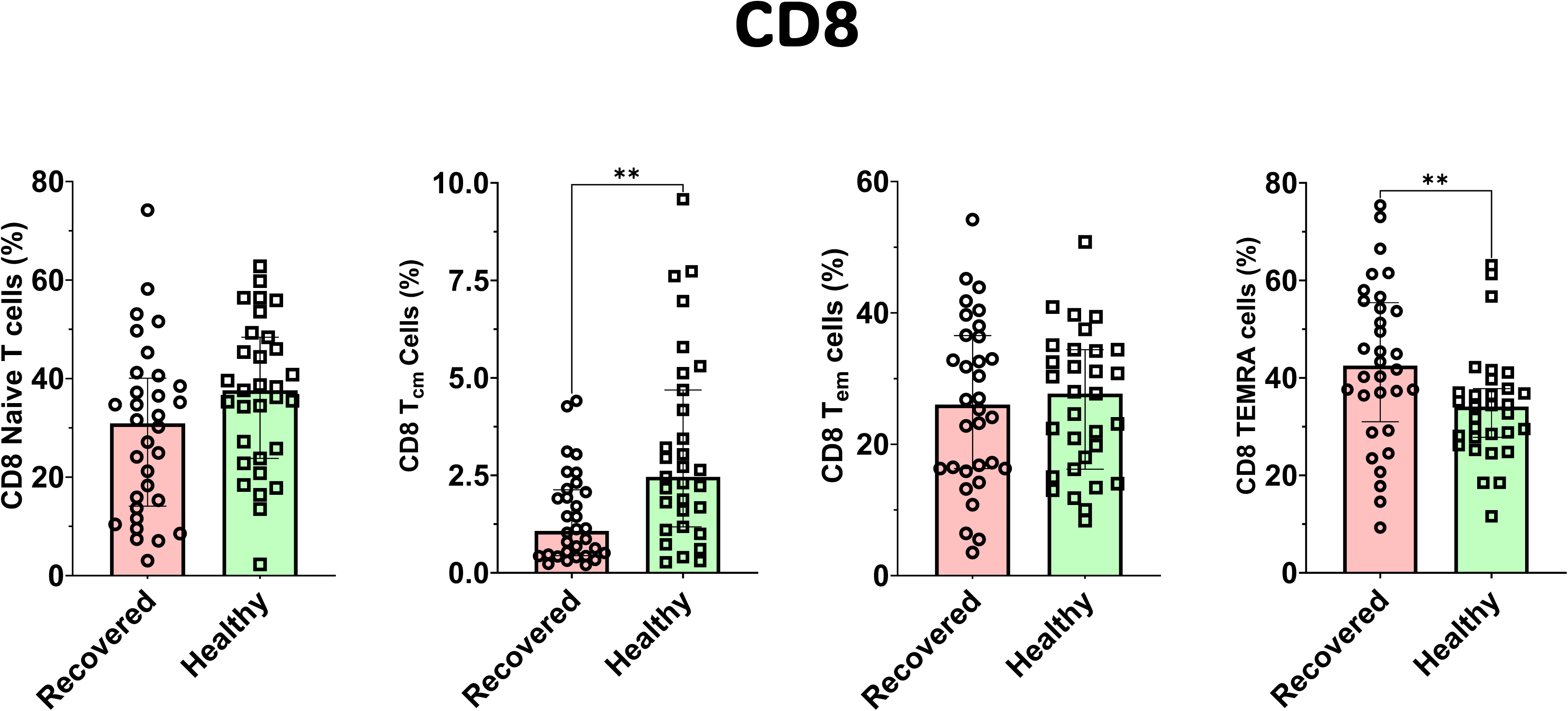
Distribution of CD8 T cell subsets (naïve, central memory, effector memory and TEMRA) among typhoid-recovered and healthy participants.

## 4. Discussion

The findings of this study provide insight into the humoral response against different classes of antigens and cellular responses against *S.* Typhi in individuals from an endemic region. We observed a robust IgM response three months post-infection against all the antigens we tested. In general, the presence of IgM antibodies suggests a recent infection while IgG antibodies imply long-term exposure. However, in certain viral and bacterial infections, long-lasting IgM response has been observed (28–30). Another possibility for a prolonged IgM antibody response is possibly due to a continued presence of the pathogen or re-exposure which is common in endemic areas. The persistence of IgM antibodies further complicates the interpretation of immune status in endemic populations (31). Interestingly, while the IgM antibody levels were elevated in case of the typhoid-recovered individuals as compared to the healthy group, IgG levels were not substantially different. These results imply that IgG mediated humoral immunity against *S.* Typhi returned to its baseline level during the post three-month recovery period. Alternatively, healthy participants likely have background IgG from asymptomatic exposures which is leading to the underestimation of the IgG levels present in recovered individuals.

Another key finding of this study is the better IgA response among typhoid-recovered individuals against polysaccharide antigens (Vi-PS and LPS). Vi capsule and LPS are the antigens that are heavily expressed on *S.* Typhi and engage with immune cells. IgA antibodies are present in the mucosal secretions and are known to contribute to the local immunity in the gut, potentially preventing *S.* Typhi from establishing the infection (9,10). IgA response against LPS was also found to be an accurate diagnostic test in a human challenge study (32). These results suggest that we may need to test the immune responses against such other antigens of *S.* Typhi to expand the repertoire of the options available to be used for multigenic screening for Typhoid.

Polysaccharides are classic T cell independent antigens and primarily trigger IgM and IgG2 responses but generate little immunological memory (33). In regions with frequent exposure, repeated encounters with antigens tend to sustain IgM levels. In contrast, immune response against protein antigens are T cell dependent (34). They usually induce IgG antibodies and promote long-term memory. In this study, IgG levels to HlyE, H protein and OMP showed no significant difference between groups after three months of disease. This could reflect the waning of IgG over time towards the basal level which is significantly high in endemic regions.

Another aspect explored in this study was the distribution of circulating B cell and T cell subsets among typhoid-recovered and healthy individuals. The overall composition of major B cell subsets was largely comparable between the two groups. However, transitional B cells showed elevated frequencies among typhoid-recovered individuals compared with healthy participants, indicating possible alterations in early B cell maturation following infection. The presence of HlyE-specific memory B cells in both groups further suggests that individuals living in endemic regions may encounter the pathogen or related antigens even in the absence of clinically diagnosed disease. Interestingly, differences were more prominent in the CD8 T cell compartment. Typhoid-recovered individuals showed higher frequencies of terminally differentiated TEMRA cells, whereas healthy individuals exhibited relatively higher proportions of central memory CD8 T cells. Elevated levels of TEMRA cells may be a result of repeated antigen exposure and are associated with cytotoxic effector function (35). Therefore, the increased frequency of this subset in recovered individuals may reflect antigen-driven differentiation during infection. These observations support the idea that immune responses to S. Typhi in endemic regions are shaped not only by visible infection but also by repeated exposure to the pathogen.

The antibody-dependent NK cell activation (ADNKA) assay showed that the plasma from typhoid-recovered individuals did not significantly enhance NK cell activation and proliferation markers (CD16, CD25 & CD69 respectively). These findings are at variance with a previous study (36), probably because of the modified protocol we followed for this study or due to the possibly exposed healthy individuals from whom PBMCs were used for this assay. The mature, CD56^dim^CD16+ NK cells are identified as a highly differentiated subset known for their potent cytotoxicity and capacity to induce antibody-dependent cellular cytotoxicity (ADCC); on the other hand, less mature CD56^bright^CD16-NK cells are major cytokine producers with regulatory functions (37). The enigmatic behavior of NK cell responses in this study could be attributed to several factors. One possibility is that the plasma from typhoid-recovered individuals possibly contains immunomodulatory factors that suppress NK cell activity, similar to what is observed in other chronic infections like Hepatitis C and HIV (38,39).

A limitation of this study is the lack of acute-phase samples, which might have given clear differentiation of the immune parameters tested between healthy and infected individuals. Another limitation is the age group of our participants which are mainly in the adult category and hence even healthy participants are probably exposed to the pathogen due to its endemicity in the region. Therefore, understanding the impact of endemicity on the immune response to typhoid with a larger sample size is essential for developing effective prevention and control strategies. Despite these limitations, this study provides critical insights into immune responses to *S.* Typhi antigens in endemic regions, offering an understanding of the immune correlates which could be protective. Moreover, the healthy population at a larger sample size may need to be screened against different antigens and antibodies to draw a cut-off to differentiate the symptomatic from exposed individuals, particularly for endemic regions. Additionally, the pre-existing immunity in healthy individuals, as reflected by their antibody levels, could be a result of prior, subclinical infections, contributing to the overall immune landscape of the population (40). Further, many of the recent vaccines against typhoid are having Vi-polysaccharide as a component, this study could provide a hindsight for the kind of immune response one can expect in an endemic region against the Vi-antigen. Together, these results emphasize the need for tailored vaccine strategies that considers pre-existing immunity and endemic exposure.

## 6. Acknowledgements

The authors wish to acknowledge BRIC THSTI, Faridabad for intramural funding to support this work (THSTI Core/EX-DBT-55-2022/P366). We are thankful to Jishnu Sankar and Dr. Dinesh Mahajan for help with biotinylation of the Vi-PS; and to Harshit Singh, Nafees Ahmed, Anmol Gaur, Sakshi Gaur, Manas Ranjan Tripathy and Jitender Chandilla, Immunobiology and Immunotherapy laboratory, BRIC THSTI, Faridabad for the suggestions and help. The authors also thank Mr. Nitesh Dhull and Dr. Sankalp for their help with the samples and ethical approval. We are particularly thankful to the participants for donating their blood samples, which made this study feasible.

## 7. Author’s contribution

Atharv Athavale collected samples, performed experiments, made tables and figures, wrote, reviewed and edited the different versions of the manuscript; Adarsh Subramaniam and Aftab Hussain-collected samples and performed experiments; Indu Rani-performed experiments; Deepak Kumar Rathore and Santosh K. Upadhyay-edited and reviewed the different versions of the manuscript; Anil Kumar Pandey and Praveen Kumar Malik provided samples for the study, reviewed the different versions of the manuscript; Amit Awasthi conceptualized the study, and reviewed the different versions of the manuscript; Ramesh Chandra Rai conceptualized the study, wrote the manuscript, reviewed and edited the different versions of the manuscript, tables and figures. All authors read and approved the final version of the manuscript.

## 9. Data Availability

Data related to the article has been presented in the form of figures and tables. Keeping confidentiality of the participants the details could be shared as per the journal’s policy.

