## Supplementary figures and images for "Immune Responses to Salmonella Typhi Antigens among Typhoid Recovered Individuals in an Endemic Region"

### Supplementary figure 1

#### Slide 1
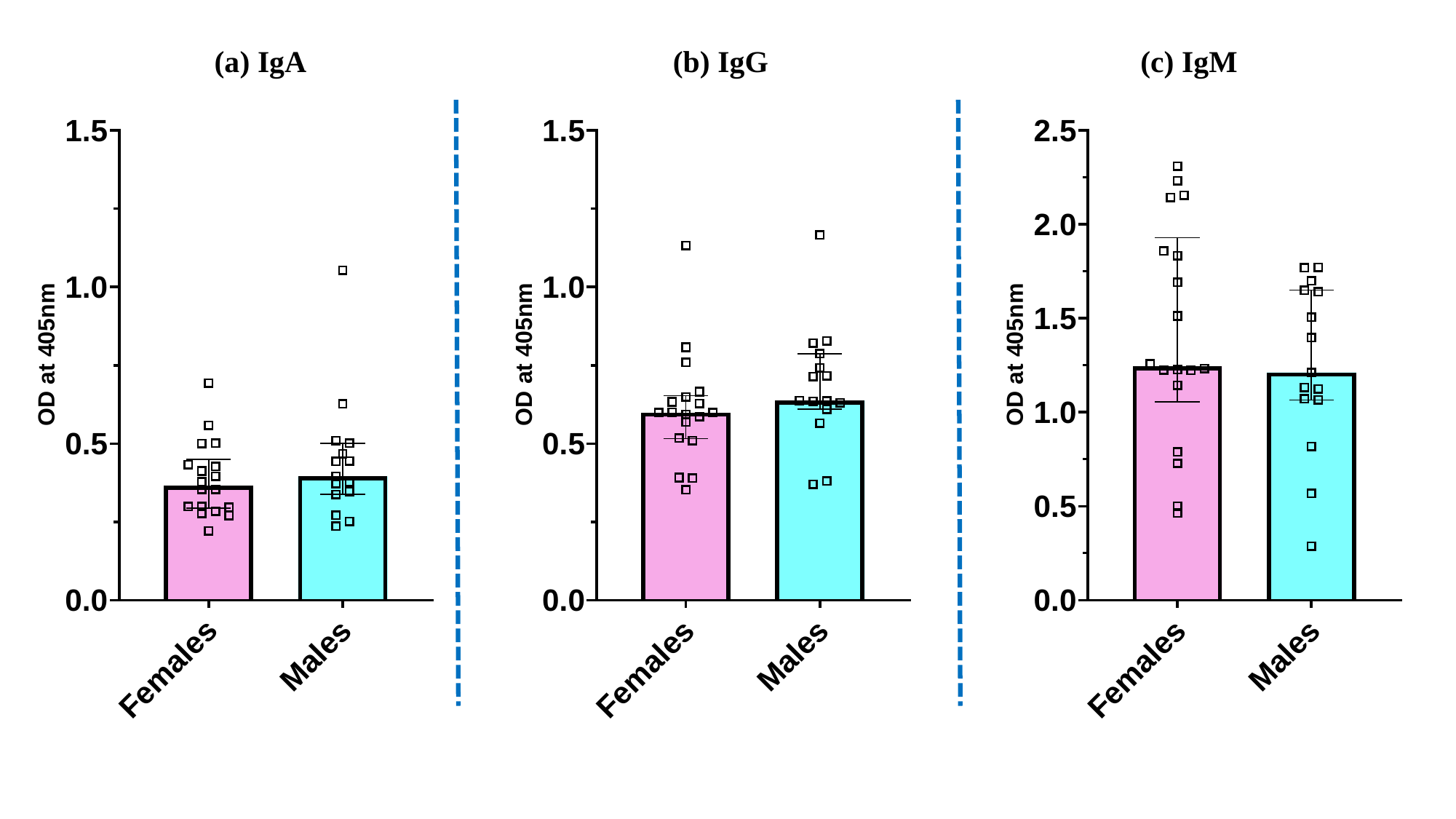

(a) IgA
(b) IgG
(c) IgM

### Supplementary figure 2

#### Slide 1
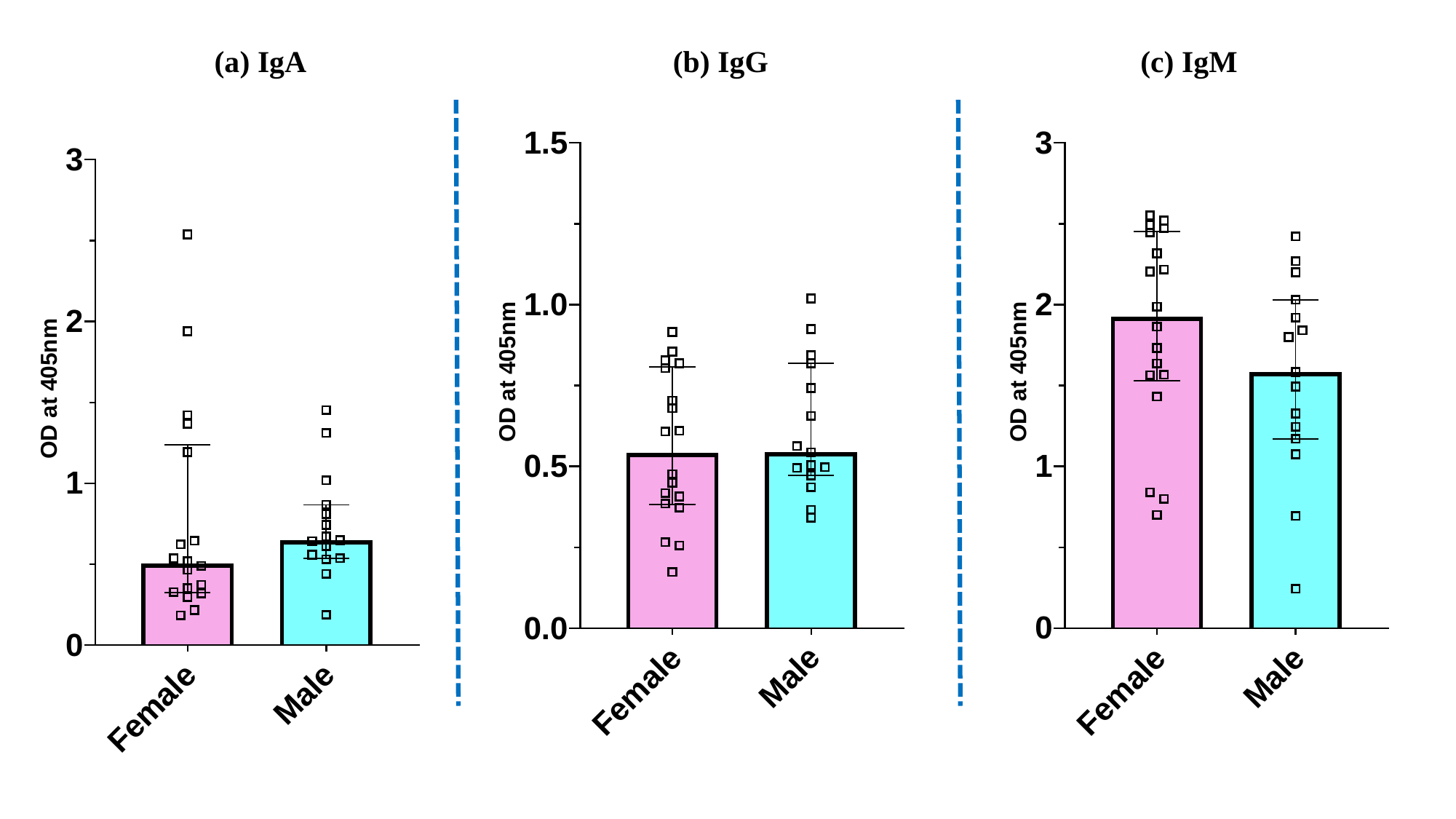

(a) IgA
(b) IgG
(c) IgM

### Supplementary figure 3

#### Slide 1
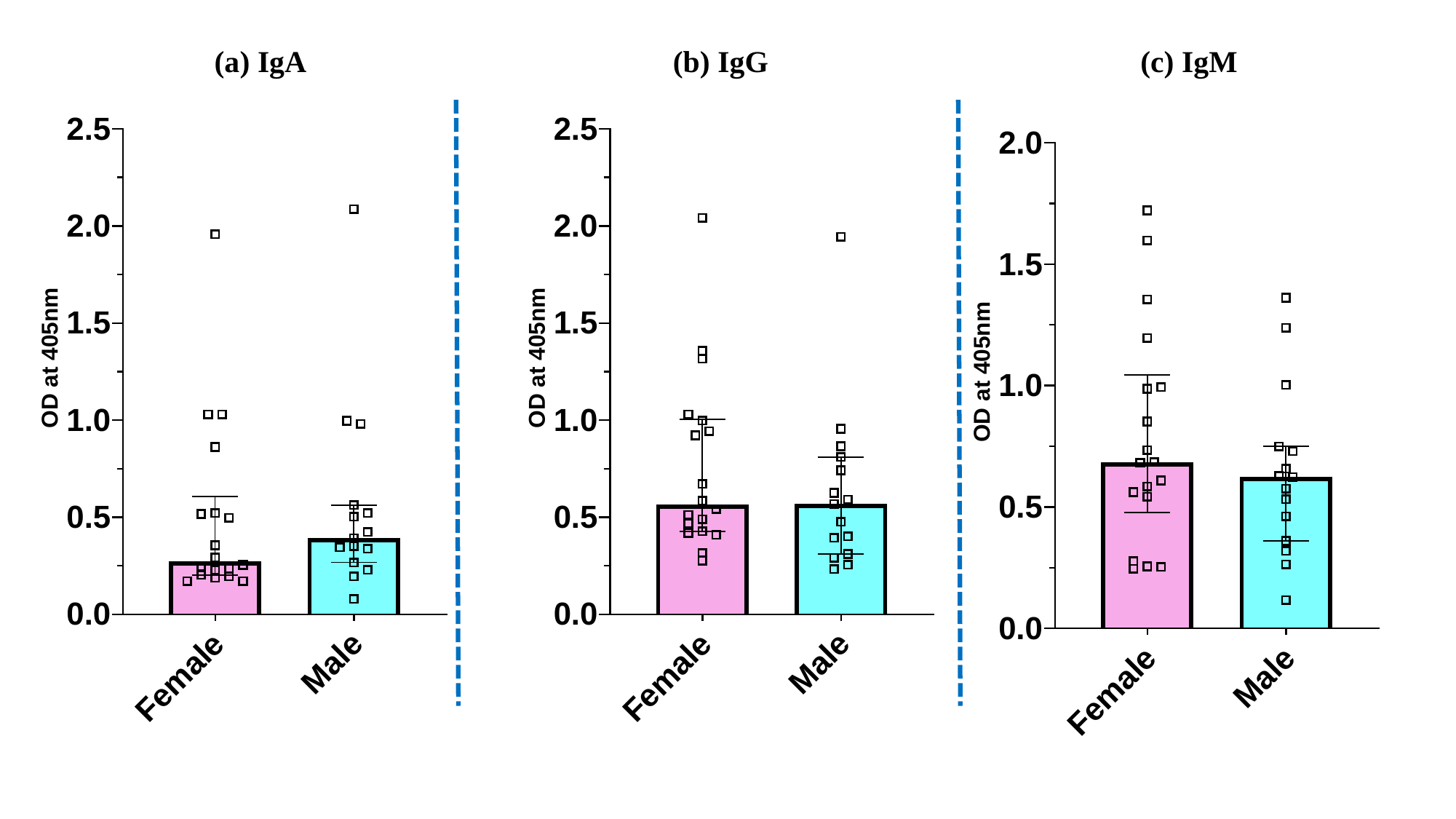

(a) IgA
(b) IgG
(c) IgM

### Supplementary figure 4

#### Slide 1
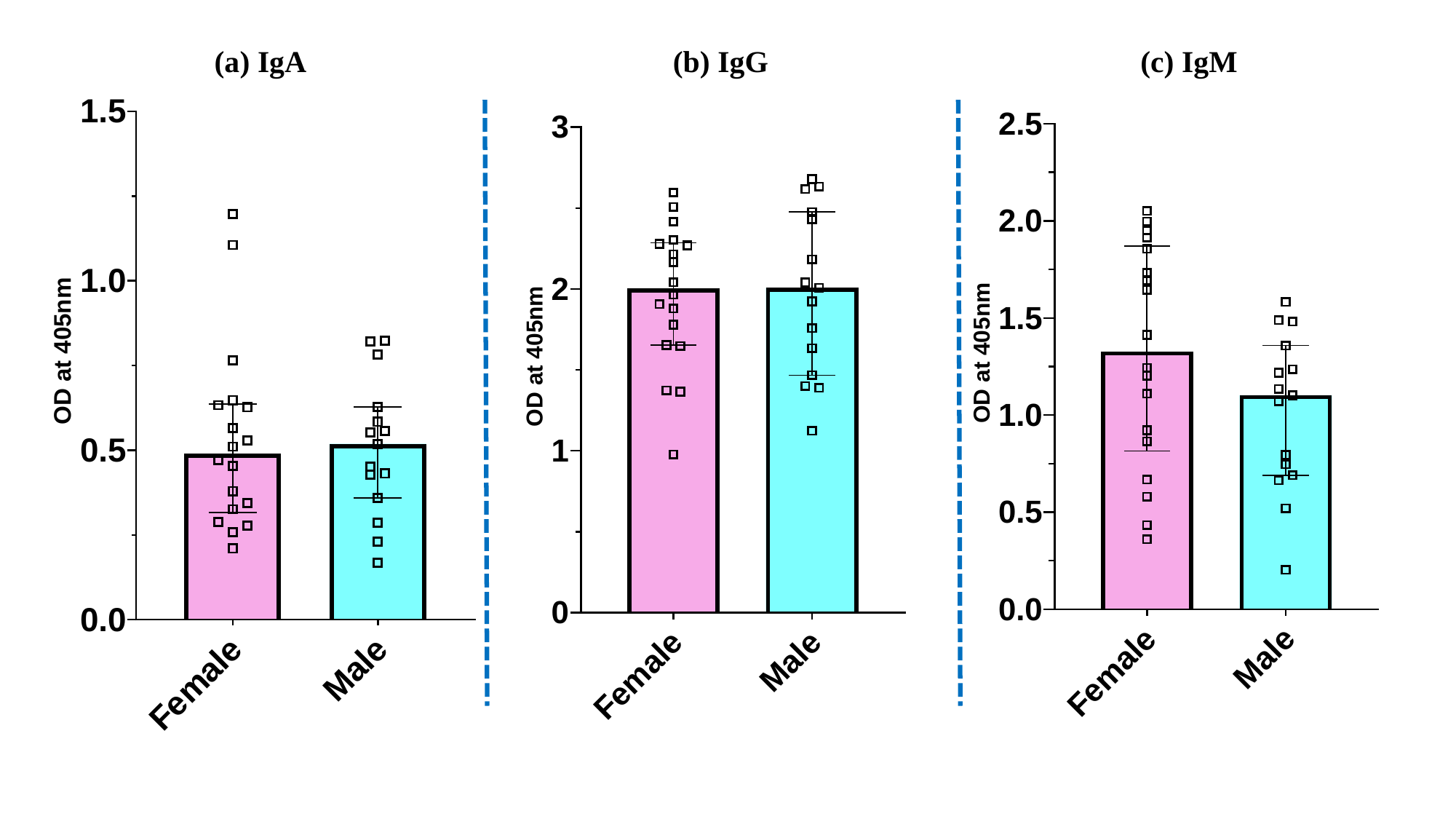

(a) IgA
(b) IgG
(c) IgM

### Supplementary figure 5

#### Slide 1
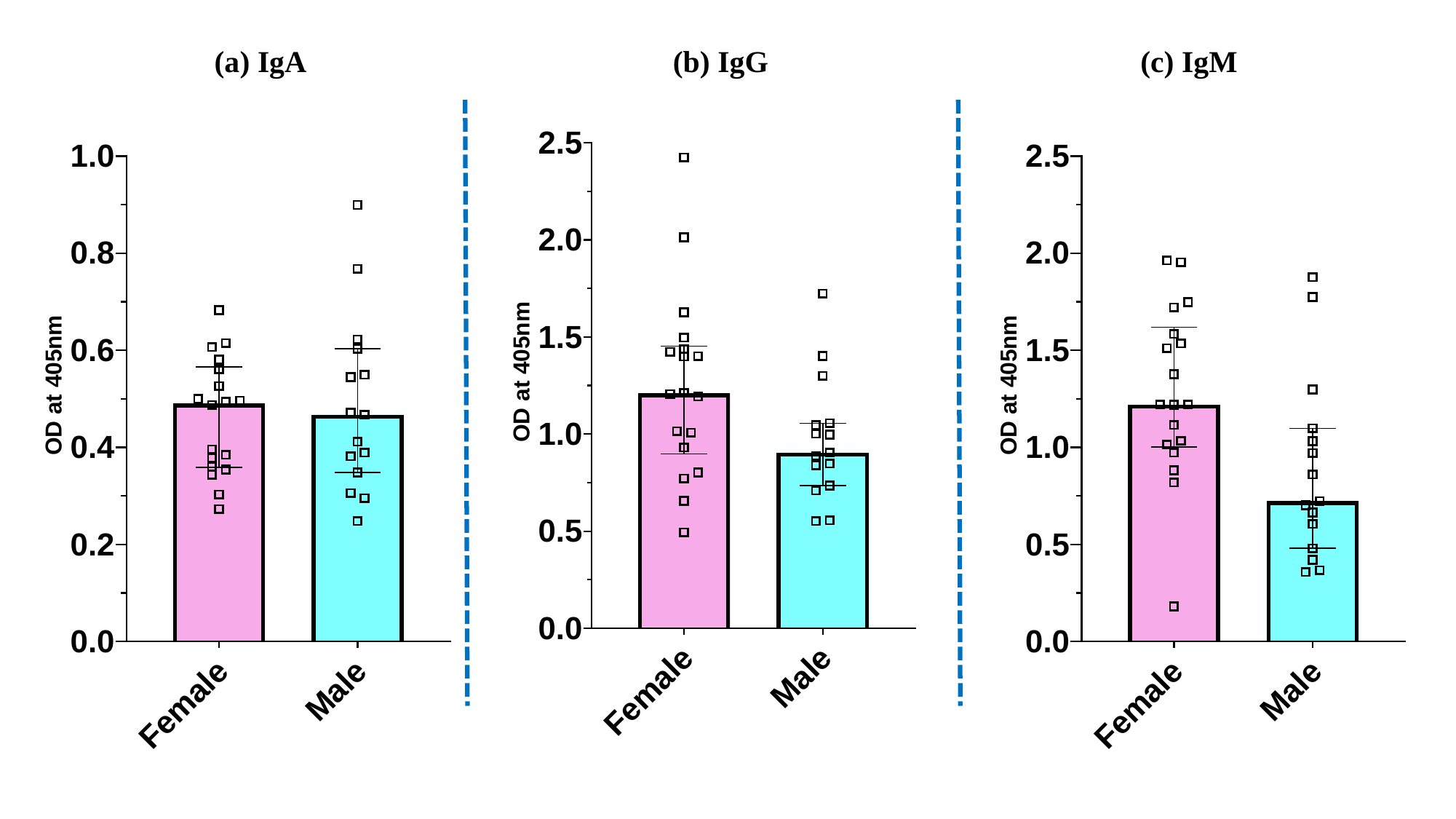

(a) IgA
(b) IgG
(c) IgM
